# Metals enable a non-enzymatic acetyl CoA pathway

**DOI:** 10.1101/235523

**Authors:** Sreejith J. Varma, Kamila B. Muchowska, Paul Chatelain, Joseph Moran

## Abstract

The evolutionary origins of carbon fixation, the biological conversion of CO_2_ to metabolites, remain unclear. Phylogenetics indicates that the AcCoA pathway, the reductive fixation of CO_2_ to acetyl and pyruvate, was a key biosynthetic route used by the Last Universal Common Ancestor (LUCA) to build its biochemistry. However, debate exists over whether CO_2_ fixation is a relatively late invention of pre-LUCA evolution or whether it dates back to prebiotic chemistry. Here we show that zero-valent forms of the transition metals known to act as co-factors in the AcCoA pathway (Fe^0^, Ni^0^, Co^0^) fix CO_2_ on their surface in a manner closely resembling the biological pathway, producing acetate and pyruvate in near mM concentrations following cleavage from the surface. The reaction is robust over a wide range of temperatures and pressures with acetate and pyruvate constituting the major products in solution at 1 bar of CO_2_ and 30 °g;C. The discovered conditions also promote 7 of the 11 steps of the rTCA cycle and amino acid synthesis, providing a stunning direct connection between simple inorganic chemistry and ancient CO_2_-fixation pathways. The results strongly support the notion that CO_2_-fixation pathways are an outgrowth of spontaneous geochemistry.

---

From the very earliest stages of the transition from chemistry to biochemistry at the origin of life, synthetic pathways must have operated to build molecular complexity from simple starting materials.^1,2^ Most efforts to understand synthetic prebiotic chemistry have focused on synthesizing biologically relevant compounds through de novo chemical routes that bear minimal resemblance to their biosynthesis.^3^ An alternative and more parsimonious possibility is that some metabolic pathways are “chemical fossils” of primitive chemistry that operated before the emergence of enzymes, RNA or cells.^4,5^ These extant anabolic pathways might have originated as chemical paths of least resistance to relieve pent-up redox gradients between the reduced iron that formed the early Earth’s bulk and its comparatively oxidized CO_2_-rich atmosphere and oceans.^6^ The anabolic systems of greatest interest in this regard are two CO_2_-fixing pathways used by autotrophic organisms^7,8^: the reductive AcCoA pathway^9^ (also known as the Wood-Ljungdahl pathway), which converts CO_2_ to acetyl CoA and then to pyruvate, and the reductive tricarboxylic acid cycle^10^ (rTCA cycle; also known as the reverse Krebs cycle).^11,12^ Phylogenetic studies indeed point to the presence of the AcCoA pathway in the last universal common ancestor.^13^ An ancestral pathway consisting of the Ac-CoA pathway and an incomplete form of the rTCA cycle is proposed to have been operative in prebiotic chemistry and consequently in early life.^14^ Other studies suggest these two pathways were once unified in their complete forms as part of a larger network that embodies stabilized network autocatalysis, a feature that could serve to amplify its constituent chemicals (Figure 1a).^15,16^ While these conceptions of prebiotic chemistry provide strong explanatory power for why the structure of metabolism is as it is, supporting chemical evidence is critically lacking.^17,18,19^ In this context, our laboratory is engaged in systematic efforts to uncover non-enzymatic chemistry related to these biological CO_2_-fixation pathways. We recently reported that all of the reduction and (de)hydration reactions of the rTCA cycle can be driven in sequence under a common set of conditions by the combination of Zn^2+^, Cr^3+^ and Fe^0^.^20^ In this paper, we turn our attention to the search for simple chemical reagents for the first two C-C bond-forming events within a hybrid AcCoA pathway/rTCA cycle; the reductive fixation of two molecules of CO_2_ to generate an acetyl group and further reductive carboxylation to form pyruvate. These metabolic reactions rely heavily on catalysis by metalloenzymes and co-factors utilizing Fe, Ni, Co, Mo or W,^21,22,23,24^ but the mechanistic steps they enable - reductions, dehydrations and migratory insertions – are also encountered in classical transition metal catalysis (Figure 1b).

**Figure 1.**
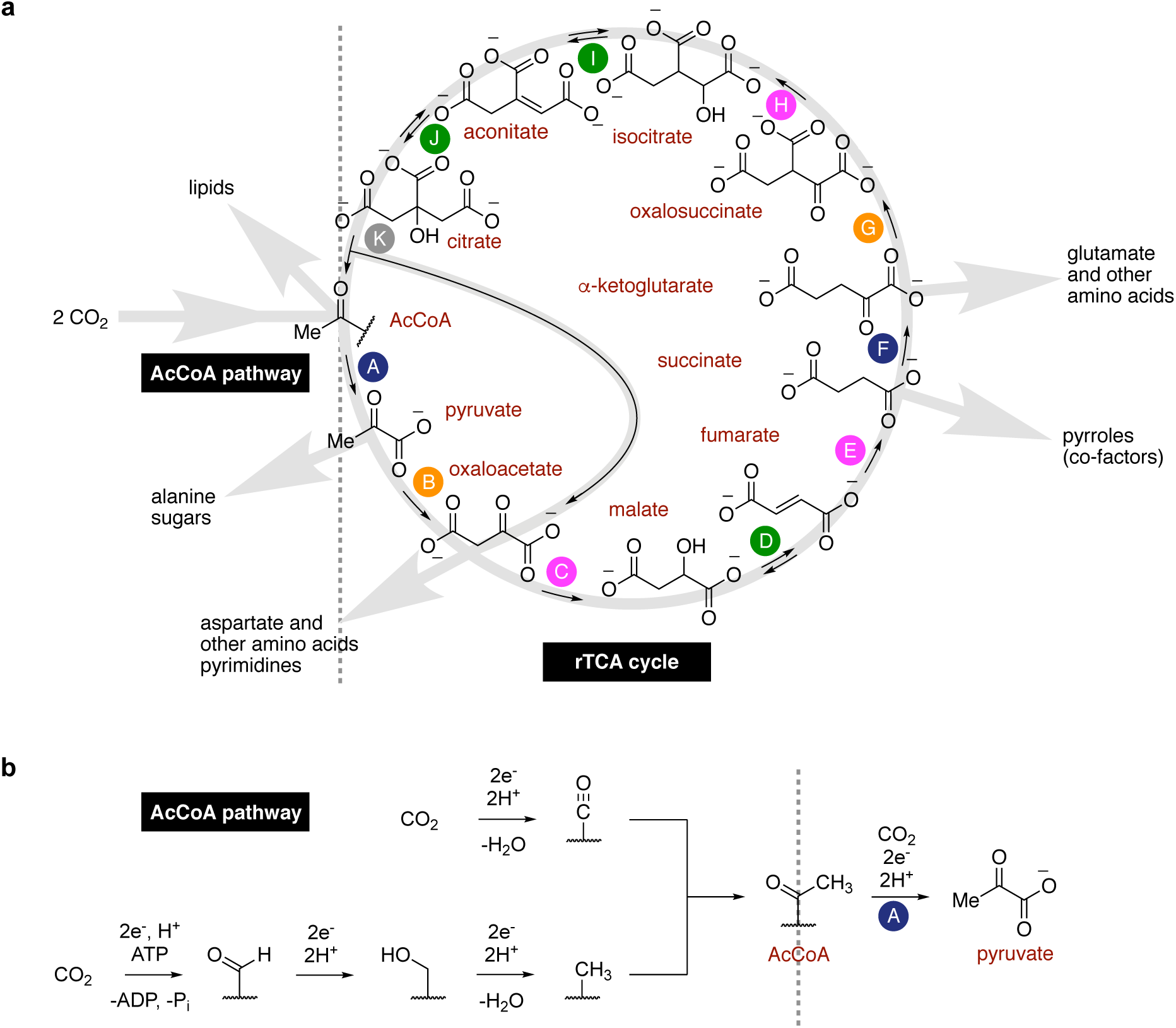
(a) Hypothetical proto-anabolic network consisting of the AcCoA pathway and the rTCA cycle, showing the role of its intermediates as universal biosynthetic precursors. (b) Mechanistic outline of the AcCoA pathway and the first reductive carboxylation (step A) of the rTCA cycle, giving pyruvate. For clarity, organic and metallic co-factors are depicted as a squiggly line.

## Results and discussion

Anticipating a link between the role of metallic cofactors in extant microbial metabolism and in prebiotic chemistry,^25,26,27^ we set out to investigate whether simple zero-valent forms of the metals involved in the reductive AcCoA pathway and in pyruvate biosynthesis might promote C-C bond formation from CO_2_ in water. We initially screened reactions of 1 mmol of Fe, Co, Ni, Mn, Mo and W powders (for specifications see Table S1) in a 1 M KCl solution in deionized H_2_O by heating to 100 °C under 35 bar CO_2_ pressure for 16 h. The reaction mixtures were then treated with KOH to precipitate hydroxides, which were removed by centrifugation prior to analysis by ^1^H NMR and GC-MS (Figures S1 – S8). Quantification was achieved by comparison to a calibration curve prepared from authentic samples (Figure S9). To our surprise, all of these metals were found to promote the formation of acetate in up to 0.28 ± 0.01 mM concentration, with considerable amounts of pyruvate observed in the cases of iron (0.03 ± 0.01 mM), nickel (0.02 ± 0.00 mM) and cobalt (0.01 ± 0.00 mM) (Figure 2). Substantial quantities of formate (up to 2.3 ± 0.2 mM in the case of cobalt) and methanol (up to 0.39 ± 0.00 mM in the case of molybdenum) were also found in almost all cases. Control experiments in the absence of metal powders or in the absence of CO_2_ did not produce detectable quantities of carbon fixation products (Figures S10 and S11).

**Figure 2.**
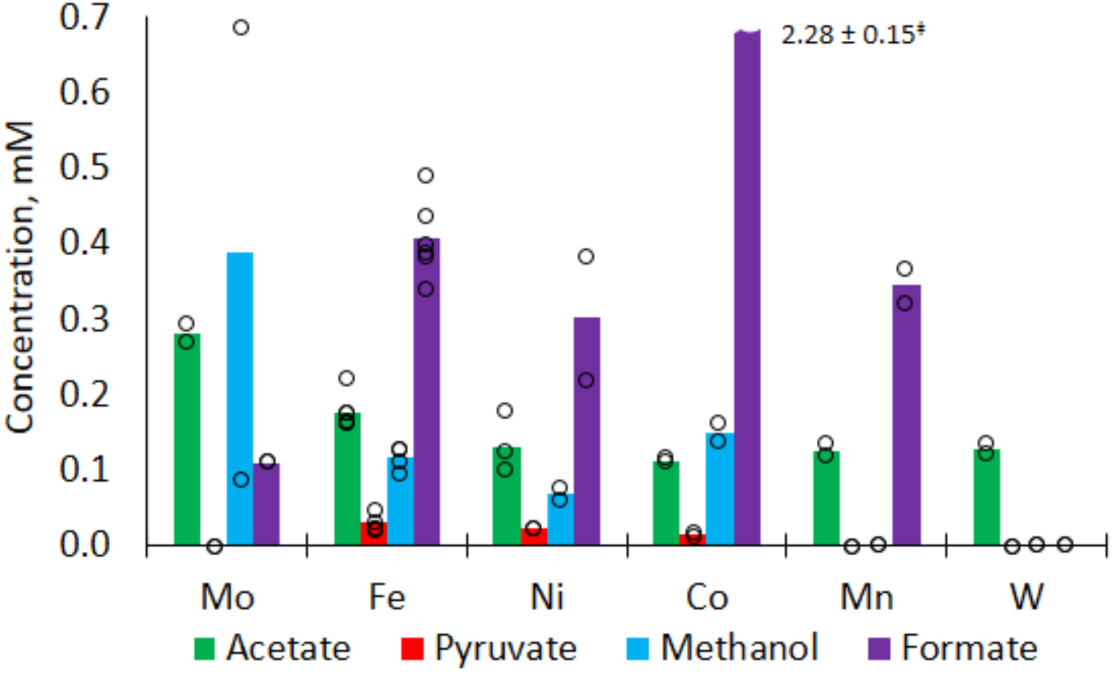
Carbon fixation by metals under mild, hydrothermal conditions: 100 °C, 35 bar CO_2_, 1 M KCl in H_2_O, pH = 7 (except for Mo, where the initial unbuffered pH was 2), 16 h (see SI for experimental and analytical details, as well as error analysis). Bar chart shows mean values of at least two independent runs. *‡* Formate concentrations: 2.13 mM, 2.44 mM; reported error corresponds to mean standard deviation.

In light of iron’s position as the premier Earth-abundant metal^28^ and its demonstrated ability to drive non-enzymatic reduction reactions of the rTCA cycle,^20^ we elected to study iron-mediated CO_2_ fixation in more detail by evaluating the influence of temperature, pressure, time, pH, salt identity and salt concentration on the reaction outcome. First, we studied the effect of temperature over the range 30-150 °C under typical conditions (35 bar CO_2_, unbuffered 1 M KCl solution in deionized H_2_O, 16 h) (Figure 3a, see also Figure S12 and Table S2). At the lower end of the temperature range, acetate is the major product in solution, with pyruvate and formate formed in slightly smaller quantities. Increasing the reaction temperature to 100 °C results in the appearance of methanol among the other products. At 150 °C, pyruvate is no longer observed, presumably due to thermal decomposition. The reaction is robust to CO_2_ pressure over the investigated range 1-40 atm at 30 oC, with acetate and pyruvate being the major products at lower pressures (Figure 3b, see also Figure S13 and Table S3). The reaction progress was monitored at different times under two representative sets of conditions. At 30 °C and 1 bar CO_2_ acetate and pyruvate are the major products, with the former reaching nearly 0.7 mM after 40 h before decreasing in concentration (Figure 3c). At 100 °C and 35 bar CO_2_ the initial buildup of formate is rapid, increasing to 7.2 ± 0.4 mM after 6 h before decreasing sharply as acetate and pyruvate begin to appear (Figure 3d, see also Figure S14 and Table S4). After 60 h, acetate and pyruvate reach maximal concentrations of 1.09 ± 0.00 mM and 0.1 ± 0.01 mM, respectively. Stopping the reaction at this time reveals that no Fe^0^ visibly remains, though only ≈1% of the available electrons from Fe^0^ were channeled towards the described C1-C3 products in solution (assuming Fe^2+^ is the terminal product, see Table S5), indicating that Fe^0^ predominantly reacts to form gaseous products under these conditions.^29^ By 85 h, the concentrations of all products in solution decrease, and ethanol is detected in the reaction mixtures for the first time (Figure S15). The disappearance of the Fe^0^ “fuel” necessary to maintain the reaction in a far-from-equilibrium steady state presumably causes the thermal decomposition of C2 and C3 products to outcompete their generation.^30^ Neither changing the ing the initial unbuffered pH nor swapping the K^+^ electrolyte for other biologically relevant inorganic cations (Na^+^, Mg^2+^ and Ca^2+^) had significant influence on carbon fixation (Figures S16 – S26 and Tables S6 – S17). However, the salt concentration had a more significant effect, with a drop of acetate and pyruvate yields by over 20% in the absence of KCl (Figure S27 and Table S18).

**Figure 3.**
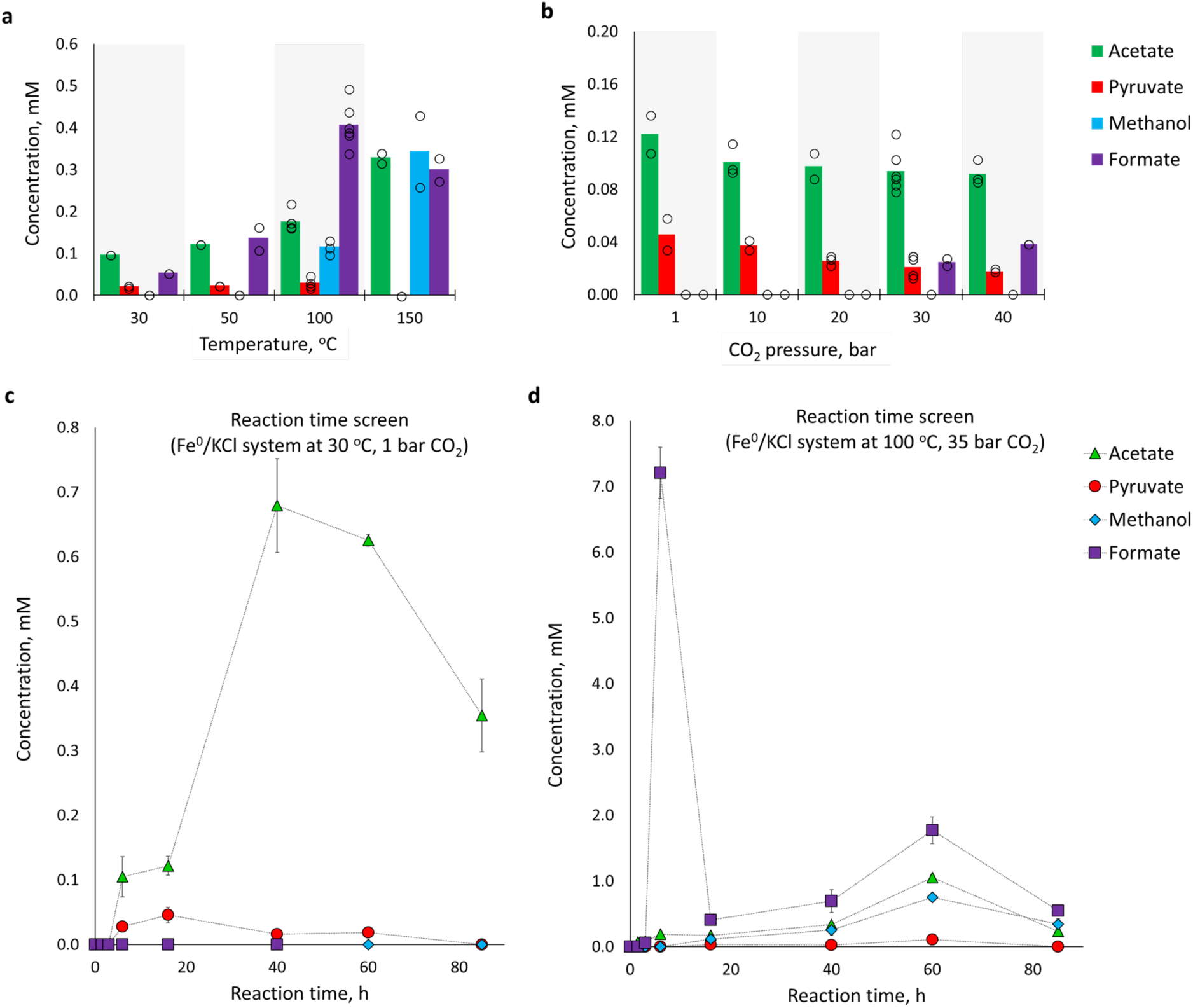
Effect of temperature, pressure and reaction time on iron-promoted CO_2_ fixation in aqueous solution. **a** Effect of temperature (35 bar CO_2_; 16 h). **b** Effect of CO_2_ pressure (30 °C; 16 h). **c** Reaction progress over time (30 °C; 1 bar CO_2_). **d** Reaction progress over time (100 °C; 35 bar CO2). All reactions are 1 mmol Fe in 1 mL of a 1 M KCl solution. In **a** and **b** bar charts show mean values of at least two independent runs. In **c** and **d** error bars correspond to the mean average deviation from at least two independent runs. Lines connecting the data points do not represent a model fit.

Several additional experimental observations helped to gain insight into the mechanism of the reaction. First, in the absence of basic workup with KOH prior to NMR analysis, no C1 – C3 carbon fixation products were observed in solution (Figure S29). Second, introduction of formate, methanol or acetate into the reactor under typical reaction conditions did not result in their conversion to higher C2 or C3 products (Figure S30). Thus, formate, methanol and acetate, free in solution, do not appear to be intermediates in the reaction. Third, lactate, the product of parasitic reduction of pyruvate, was never observed in any of our experiments, despite the fact that it is readily detected upon exposure of an aqueous pyruvate solution to metallic Fe (Figure S30d). Interestingly, Fe-mediated carbon fixation reactions performed with CO in-stead of CO_2_ at 30 °C produced only tiny amounts of acetate and no detected pyruvate (Figure S31 and Table S19). On the basis of these observations and the reaction’s kinetic profile, we propose a preliminary mechanism whereby carbon fixation °Ccurs on the surface of the reduced metal to produce surface-bound species. Initial reduction of CO_2_ °Ccurs to generate surface-bound carbon monoxide and formyl groups. Further reductions of the formyl group with expulsion of water leads to a surface-bound methyl group. Chain growth via migratory insertion of carbon monoxide into the methyl group produces an acetyl species, which itself can undergo further migratory insertion of CO to furnish a surface-bound pyruvyl species. The resistance of the pyruvyl species to reduction by Fe^0^ may be rationalized by a diminished reactivity of the surface-bound species compared to pyruvate in solution. Basic workup at the end of the reaction with KOH is therefore required to cleave the surface-bound formyl, methyl, acetyl or pyruvyl species to furnish formate, methanol, acetate or pyruvate in solution, respectively (Figure 4). The kinetic profile observed in Figure 3c may therefore be interpreted as an initial saturation of the Fe surface with formyl and CO groups, followed by subsequent reactions of those groups to produce acetyl and, eventually, pyruvyl groups. Further study is required to distinguish between the intermediacy of surface-bound metal carboxylates and surface acyl metal species. Regardless, the proposed mechanistic picture is reminiscent of the enzymatic mechanisms of the reductive AcCoA pathway and the first step of the rTCA cycle.^27^

**Figure 4.**
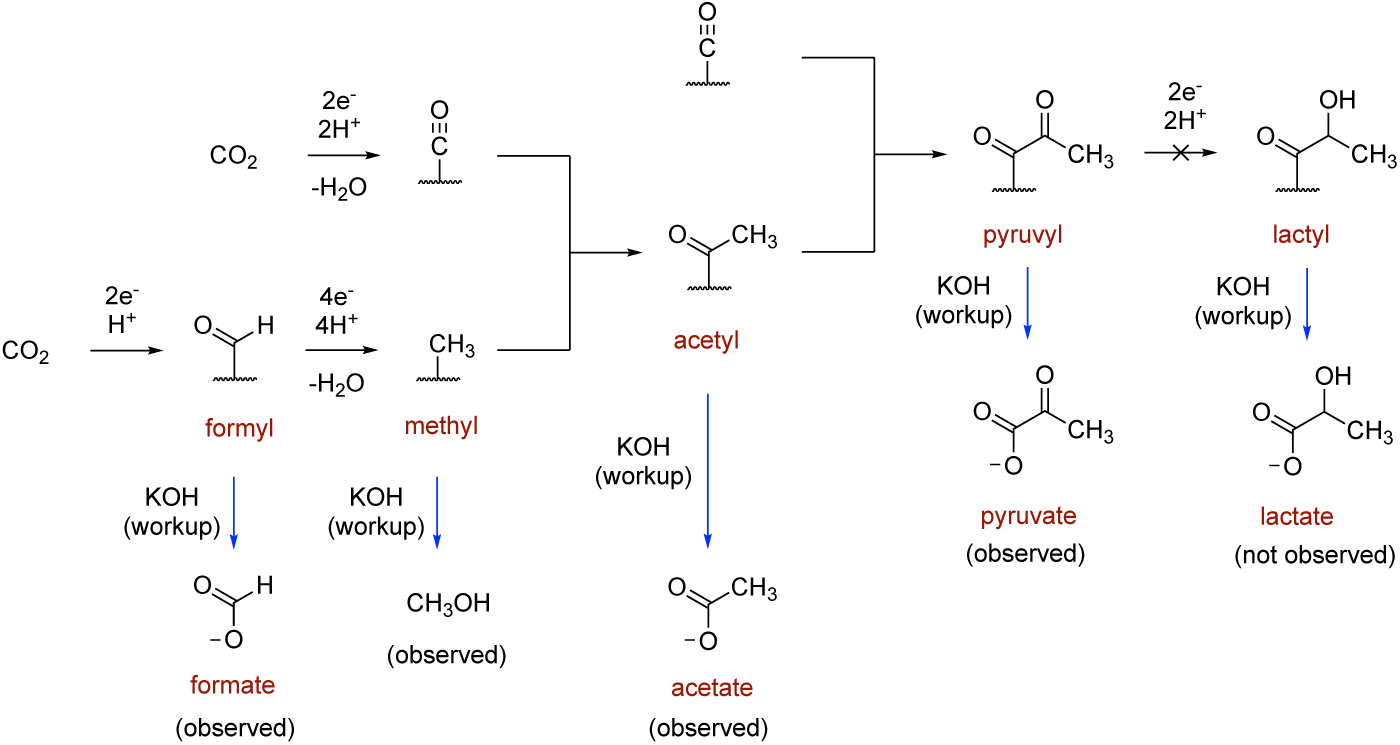
Plausible mechanism for carbon fixation on the surface of Fe^0^ and the detection of formate, methanol, acetate and pyruvate in aqueous solution following hydrolysis with KOH. The depicted surface bound structures may represent a surface-bound carboxylate or an acyl metal species.

Finally, we set out to test the mutual compatibility of the carbon fixation conditions with other steps of the rTCA cycle. In our previous work, Fe^0^, Zn^2+^ and Cr^3+^ was found to promote the three-step oxaloacetate-to-succinate sequence and the three-step oxalosuccinate-to-citrate sequence (Figure 1a, steps C-E and steps H-J), however, strongly acidic conditions were required (1 M HCl in H_2_O).^20^ When oxaloacetate was heated for 16 h in the presence of Fe powder and Cr^3+^ at 140 °C under 35 bar of CO_2_, an appreciable amount of succinate was formed alongside the carbon fixation products, as evidenced by NMR and GC-MS (Figures S32 and S33). An analogous experiment carried out with *in situ* generated oxalosuccinate (heated at 140 °C for 16 h under 35 bar of CO_2_, in the presence of Fe powder and Cr^3+^) resulted in the formation of citrate (detected by GC-MS, Figure S34). Although both of these reaction sequences are less selective under the mildly acidic conditions afforded by CO_2_-saturated water than in 1 M HCl, they nonetheless suggest the potential compatibility of the carbon fixation conditions shown here with the non-enzymatic promotion of rTCA cycle reaction sequences. Lastly, the introduction of hydrazine into the otherwise identical Fe-rich conditions allowed for the non-enzymatic synthesis of the amino acid alanine from pyruvate (Figure S35).^20^

## Conclusions

We have shown that zero-valent forms of metals used by metalloenzymes to catalyze biological carbon fixing pathways are able to promote CO_2_ fixation resembling the AcCoA pathway and the first step of the rTCA cycle in an experimentally trivial manner that is robust to changes in temperature, pressure, salts and pH. Acetate is produced as the major C2 product for all metals studied, whereas Fe^0^, Ni^0^ and Co^0^ produce pyruvate in up to ∼0.1 mM concentrations. The reaction operates even at 30 °C and 1 atm CO_2_ pressure and is highly selective for acetate and pyruvate under these conditions, offering a striking experimental parallel between simple chemistry and chemoautotrophic CO_2_-fixation. Indeed, CO_2_-fixing organisms using Fe^0^ corrosion as their sole source of electrons still exist on the Earth today.^31,32^ Control and time course experiments support a mechanism where CO_2_ fixation °Ccurs on the surface to produce surface-bound intermediates, notably reminiscent of some aspects of Wächtershäuser’s initial theory of surface metabolism.^4^ The temperature limits and critical need to liberate surface-bound species with KOH prior to analysis may explain why pyruvate was not observed in previous studies on the reaction of CO_2_ with Fe^0^.^29,33^ It is also more geochemically plausible than hydrothermal syntheses relying on other C1 feedstocks, such as the synthesis of activated acetate from CO and CH_3_SH,^34,35^ the low yielding synthesis of pyruvate (0.07%) in neat formic acid at high temperature and high pressure (250 °C, 2000 bar),^36^ or using highly reducing (-1.1 V) electrochemistry on greigite electrodes.^37^ Iron is the most abundant transition metal on Earth and in meteorites.^28,38^ Fe^0^ is produced in small amounts by serpentinization,^39^ is found in the Earth’s crust as telluric iron,^40^ is produced transiently in the mantle,^41^ and is the major component of the Earth’s core. Thus, numerous plausible geological scenarios might be imagined in which Fe^0^ and other reduced metals could have been continuously produced and consumed by proto-metabolism on the early Earth. The Fe^0^-promoted carbon fixation conditions are mutually compatible with other non-enzymatic transformations, describing a reaction network equivalent to the entire reductive AcCoA pathway, 7 of the 11 reactions of the rTCA cycle and amino acid synthesis. The observation that surface-bound pyruvate is not reduced to lactate in the presence of Fe^0^ - though not yet fully understood - opens up new mechanistic possibilities for how further non-enzymatic anabolic reactions might be promoted in the face of potential parasitic off-cycle reactions.^42^ The observed reactivity represents a direct parallel between prebiotic chemistry and the CO_2_-fixing pathways used by primitive autotrophic life, strongly supporting the hypothesis that metabolism originated as prebiotic geochemistry.

## Data availability

The authors declare that the data supporting the findings of this study are available within the paper’s Supplementary Information files.

## Acknowledgments

This project has received funding from the European Research Council (ERC) under the European Union's Horizon 2020 research and innovation program (grant agreement n° 639170). Further funding was provided by a grant from LabEx “Chemistry of Complex Systems”. Dr Lionel Allouche, Dr Maurice Coppe and Dr Bruno Vincent (Institute of Chemistry, University of Strasbourg, France) are gratefully acknowledged for their assistance with NMR experiments.

## Additional information

Supplementary information is available for this paper.

Correspondence should be addressed to J.M.

## Competing financial interests

The authors declare no competing financial interests.

## References

1 Ruiz-Mirazo, K., Briones, C. & de la Escosura, A. Prebiotic systems shemistry: new perspectives for the origins of life. Chem. Rev. 114, 285–366 (2014).

2 Peretó, J. Out of fuzzy chemistry: from prebiotic chemistry to metabolic networks. Chem. Soc. Rev. 41, 5394–5403 (2012).

3 Sutherland, J. D. Studies on the origin of life - the end of the beginning. Nat. Rev. Chem. 1, 1 (2017)

4 Wächtershauser, G. Before enzymes and templates: theory of surface metabolism. Microbiol. Rev. 52, 452–484. (1988).

5 Smith, E. & Morowitz, H. J. The Origin and Nature of Life on Earth: The Emergence of the Fourth Geosphere (Cambridge Univ. Press, Cambridge, 2016).

6 Russell, M. J., Hall, A. J. & Mellersh, A. R. in Natural and Laboratory Simulated Thermal Geochemical Processes 325–388 (Springer Netherlands, Dordrecht, 2003).

7 Berg, et al. Autotrophic carbon fixation in archaea. Nat. Rev. Microbiol. 8, 447–460 (2010).

8 Hügler, M. & Sievert, S. M. Beyond the Calvin cycle: autotrophic carbon fixation in the ocean. Annu. Rev. Mar. Sci. 3, 261–289 (2011).

9 Ljundahl, L. G., Irion, E. & Wood, H. G. Total synthesis of acetate from CO2. I. Co-methylcobyric acid and co-(methyl)-5-methoxy-benzimidizolycobamide as intermediates with Clostridium thermoaceticum. Biochemistry 4, 2771–2779 (1965).

10 Evans, M. C. W., Buchanan, B. B. & Arnon, D. I. A new ferredoxindependent carbon reduction cycle in a photosynthetic bacterium. Proc. Natl Acad. Sci. USA 55, 928–934 (1966).

11 Morowitz, H. J., Kostelnik, J. D., Yang, J. & Cody, G. D. The origin of intermediary metabolism. Proc. Natl Acad. Sci. USA 97, 7704–7708 (2000).

12 Smith, E. & Morowitz, H. J. Universality in intermediary metabolism. Proc. Natl Acad. Sci. USA 101, 13168–13173 (2004).

13 Weiss, M. C. et al. The physiology and habitat of the last universal common ancestor. Nat. Microbiol. 1, 16116 (2016).

14 Martin, W., Russell, M. J., On the origin of biochemistry at an alkaline hydrothermal vent. Phil. Trans. R. Soc. B 362, 1887–1926 (2007).

15 Braakman, R. & Smith, E. The emergence and early evolution of biological carbon-fixation. PLoS Comp. Biol. 8, e1002455 (2012).

16 Braakman, R. & Smith, E. The compositional and evolutionary logic of metabolism. Phys. Biol. 10, 011001 (2013).

17 Cody, G. D. et. al. Geochemical roots of autotrophic carbon fixation: hydrothermal experiments in the system citric acid, H_2_O-(±FeS)-(±NiS). Geochemica et Cosmochemica Acta 65, 3557–3576 (2001).

18 Zhang, X. V.; Martin, S. T. Driving parts of Krebs cycle in reverse through mineral photochemistry. J. Am. Chem. Soc. 128, 16032–16033 (2006).

19 Herschy, B.; Whicher, A.; Camprubi, E.; Watson, C.; Dartnell, L.; Ward, J.; Evans, J. R. G.; Lane, N. An Origin-of-Life Reactor to Simulate Alkaline Hydrothermal Vents. Journal of Molecular Evolution. 79, 213–227 (2014).

20 Muchowska, K., et al. Nat. Eco. Evo., 10.1038/s41559-017-0311-7 (2017).

21 Can, M., Armstrong, F. A., & Ragsdale, S. W. Structure, function, and mechanism of the nickel metalloenzymes, CO dehydrogenase and acetyl-CoA synthase. Chem. Rev. 114, 4149–4174 (2014).

22 Schwarz G., Mendel R. R. & Ribbe M. W. Molybdenum cofactors, enzymes and pathways. Nature 460, 839–847 (2009).

23 Schuchmann, K. & Müller, V., Autotrophy at the thermodynamic limit of life: a model for energy conservation in acetogenic bacteria. Nat. Rev. Microbiol. 12, 809–821 (2014).

24 Furdui, C., & Ragsdale, S. W. The role of pyruvate ferredoxin oxidoreductase in pyruvate synthesis during autotrophic growth by the Wood-Ljungdahl pathway. J. Biol. Chem. 275, 28494–28499 (2000).

25 Sousa, F., & Martin, W. F. Biochemical fossils of the ancient transition from geoenergetics to bioenergetics in prokaryotic one carbon metabolism. Biochim. Biophys. Acta 1837, 964–981 (2014).

26 Morowitz, H. J., Srinivasan, V., & Smith, E. Biol. Bull. 219, 1–6 (2010).

27 Camprubi, E., Jordan, S. F., Vasiliadou, R., & Lane, N. Iron catalysis at the origin of life, IUBMB Life 69, 373–381 (2017).

28 Moore, E. K., Jelen, B. I., Giovanelli, D., Raanan, H. & Falkowski, P. G. Metal availability and the expanding network of microbial metabolisms in the Archaean eon. Nat. Geosci. 10, 629–636 (2017).

29 Guan, G. et al., Reduction of aqueous CO_2_ at ambient temperature using zero-valent iron-based composites. Green Chemistry 5, 630–634 (2003).

30 Boekhoven, J., Hendriksen, W. E., Koper, G. J. M., Eelkema, R. & van Esch, J. H. Transient assembly of active materials fueled by a chemical reaction. Science 349, 1075–1079 (2015)

31 Enning, D., Garrelfs, J. Corrosion of Iron by sulfate-reducing bacteria: New views of an old problem. Appl. Environ. Microbio. 80, 1226–1236 (2014).

32 Daniels, L., Belay, N., Rajagopal, B. S., Weimer, P. J. Bacterial methanogenesis and growth from CO_2_ with elemental Iron as the sole source of electrons. Science 237, 509–511 (1987).

33 He, C., Tian, G., Liu, Z. & Feng, S. A mild hydrothermal route to fix carbon dioxide to simple carboxylic acids. Org. Lett. 12, 649–651 (2010).

34 Huber, C. & Wächtershauser, G. Activated acetic acid by carbon fixation on (Fe,Ni)S under primordial conditions. Science 276, 245–247 (1997).

35 Chandra et al. The Abiotic Chemistry of Thiolated Acetate Derivatives and the Origin of Life. Sci. Rep. 6, 29883 (2016).

36 Cody, G. D., et. al. Primordial carbonylated iron-sulfur compounds and the synthesis of pyruvate. Science 289, 1337–1340 (2000).

37 Roldan, A. et al. Bio-inspired CO_2_ conversion by iron sulfide catalysts under sustainable conditions. Chem. Commun. 51, 7501–7504 (2015).

38 Benedix, G. K., Haack, H. & McCoy, T. J., in Treatise on Geochemistry (Second Edition), 267–285 (Elsevier, Oxford, 2014).

39 McCollom, T. M. Abiotic methane formation during experimental serpentinization of olivine. Proc. Natl Acad. Sci. USA 113, 13965–13970 (2016).

40 Klöck, W., Palme, H. & Tobschall, H. J. Trace elements in natural metallic iron from Disko Island, Greenland. Contributions to Mineralogy and Petrology 93, 273–282 (1986).

41 Frost, D. J., et. al. Experimental evidence for the existence of iron-rich metal in the earth's lower mantle. Nature 428, 409–412 (2004).

42 Orgel, L. E. The implausibility of metabolic cycles on the prebiotic earth. PLoS Biol. 6, e18 (2008).

